# On the Impact of Interhemispheric White Matter: Age, Executive Functioning, and Dedifferentiation in the Frontal Lobes

**DOI:** 10.1101/226845

**Authors:** Abigail B. Waters, Kayle S. Sawyer, David A. Gansler

## Abstract

**Introduction:** In middle age, declines in executive functioning (EF) are associated with decrements in the quality and/or quantity of white and grey matter. Recruitment of homologous regions has been identified as a compensatory mechanism for cognitive decline in later middle age, however research into neural substrates of EF has yet to be guided by dedifferentiation models. We hypothesized that frontal-parietal grey matter volume, interhemispheric white matter and intrahemispheric white matter fractional anisotropy (FA) will be predictive of EF. Further, we hypothesized that the comparative association between interhemispheric white matter and EF will increase with age, because of compensatory recruitment.

**Methods:** Neurocognitive test data, DTI, and T1 MPRAGE scans (*n* = 444) were obtained from the NKI-Rockland Sample. Structural equation modeling was used to examine the relationship between age, EF, interhemispheric white matter (forceps minor; FM), intrahemispheric white matter (superior longitudinal fasciculus; SLF), and a frontal-parietal grey matter network. EF and grey matter were modelled as latent variables, with EF examined as the criterion. Additionally, a subsample of participants aged 55-85 (*n* = 168) was analyzed to examine the influence of age related compensatory mechanisms.

**Results:** There was a significant relationship between FM, grey matter, and EF, which was fully mediated by age. There was a significant relationship between SLF and EF, which was not mediated by age. For older adults, only the age-mediated pathway from FM to EF was significant.

**Discussion:** Using structural imaging data, support was found for age-related interhemispheric mechanisms of compensation, but not intrahemispheric mechanisms.

**Key points:** (1) Neural substrates of executive functioning are not static across the lifespan. (2) In older adults, white matter becomes more salient as a structural correlate of executive functioning, as recruitment needs increase. (3) While the importance of interhemispheric white matter is mediated by age, intrahemispheric recruitment remains consistent across the lifespan, and is not the primary mechanism of age-based compensation in community dwelling older adults.

## Introduction

Age-related decline in executive functioning (EF) starts earlier and at increased rates compared to other domains of cognitive functioning, like verbal ability (Deary et al., 2009; Strauss, Sherman, & Spreen, 2006) and memory (Grober, et al., 2008). These changes emerge around age 40 and are particularly important, because there are systematic relationships between EF, practical skills in daily living, and disability in elders (Cahn-Weiner, Boyle, & Malloy, 2002; Gansler, Suvak, Arean, & Alexopoulos, 2015; Johnson, Lui, & Yaffe, 2007; Mirelman et al., 2012; Yogev-Seligmann, Hausdorff, & Giladi, 2008). As the proportion of US adults aged 65 and older is expected to increase from 14.5% in 2014 to 21.7% by 2040 (U.S. Administration on Aging, 2016), determining behavioral and biological mechanisms and predictors for EF decline is an important tool for early detection and intervention for general cognitive and adaptive functional decline. More detailed knowledge of brain systems underlying age-related cognitive decline may inform our understanding of disability and pathological processes in aging.

In this study, we examined the underlying common cognitive factor of EF. EF is a broad term that refers to a group of cognitive processes needed for coordination and regulation of behavior and cognitive functions (Diamond, 2013). Models of EF generally fall into one of two categories: those based on an underlying common factor or those based on an array of specific factors. Often there are common task demands among any given neuropsychological tests of EF (i.e. goal maintenance, working memory), where that task is saturated by a common factor (Engle, Tuholski, Laughlin, & Conway, 1999; McCabe, Roediger, McDaniel, Balota, & Hambrick, 2010) with neural substrates in parietal and lateral prefrontal cortices (Collette et al., 2005). While specific aspects of dysfunction in EF processes account for differentiated but related observed behavioral effects in daily living, there are limits to predictive power with a divisional approach (Burgess, Alderman, Evans, Emslie, & Wilson, 1998). While there is evidence to support the argument that specific components of EF account for a diversity of behavioral effects in adaptive functioning, general EF factors may have the most predictive power (Barbey et al., 2012) and were therefore the focus of this investigation.

Declines in EF in the aging brain have been linked to age-related degradations in neuroanatomical structures. White matter (WM) and grey matter (GM) tissue deterioration related to aging is primarily seen in the prefrontal, temporal, and parietal lobes (Raz & Rodrigue, 2006). These areas are a critical basis of the neural representations of the EF domains of task coordination, planning, goal maintenance, working memory, and task switching (Daniels, Toth, & Jacoby, 2006). Specifically, degradation of anterior cingulate cortex (Botvinick, Cohen, & Carter, 2004; Van Veen, Cohen, Botvinick, Stenger, & Carter, 2001), the dorsolateral prefrontal cortex (Kane & Engle, 2002; MacPherson, Phillips, & Sala, 2002), and WM of the genu of the corpus callosum (Kochunov, et al., 2012) have all been predictive of age-related changes in EF. However, there is limited research on the comparative contribution of white and GM in executive networks in large samples of community dwelling older adults.

Some studies also suggest that the root of these age-related changes in EF is a function of either, under recruitment, or, nonselective recruitment of related brain regions (Logan, Sanders, Snyder, Morris, & Buckner, 2002). These findings suggest that because EF often involves the coordination of several brain regions, individual differences in WM may increase in influence as age increases due to the need for broader recruitment. One of the most well studied neural compensatory mechanisms observed in older adults involves increased homologous recruitment and dedifferentiation of brain regions, which seem to mitigate the effect of age-related deterioration in cortical areas and its effects on EF. In this context, dedifferentiation refers to the shift from focal to more diffuse recruitment of brain networks. Previous research has established that this process emerges around age 55 (Rypma & D’Esposito, 2000), while evidence suggests EF decline begins earlier. For younger adults (age 18-25), activation is found primarily in the left hemisphere for verbal working memory tasks, and in the right hemisphere for spatial working memory. However, older adults (age 65-75) show patterns of anterior bilateral activation for both verbal and spatial working memory, recruiting homologous brain regions in the opposite hemisphere to do the same tasks (Reuter-Lorenz et al., 2000).

These patterns of dedifferentiation are often found in areas associated with working memory in the dorsolateral prefrontal cortex, an important component of EF (Reuter-Lorenz et al., 2000). Younger adults only showed high activation in dorsolateral areas when switching between tasks, whereas older adults showed activation in these regions during both isolated tasks and when switching (DiGirolamo et al., 2001). Expanded distributions of activity may be the result of greater demands for executive control as EF becomes less automated with age. For high performing, community dwelling older adults, there may be a hierarchy of compensatory activation, where homologous regions are recruited before heterologous regions (Cabeza et al, 2002).

Although the associations between prefrontal cortex volume and EF do remain consistent throughout the lifespan (Yuan & Raz, 2014), the ability to increase recruitment in older adults is largely dependent on the integrity of WM pathways (Just, Cherkassky, Keller, & Minshew, 2004; Nordahl, et al., 2006). For older adults, the ability to activate areas bilaterally results in better performance on letter matching tasks in visual laterality studies (Reuter-Lorenz, Stanczak, & Miller, 1999). For older adults, there are associations between individual differences in integrity of these interhemispheric pathways and the degree of observed compensatory activation (Perrson et al., 2006). Because of this mechanism, interhemispheric WM (e.g., forceps major) connectivity may play a more important role in older adults compared to both GM and intrahemispheric WM (e.g., superior longitudinal fasciculus).

To examine the interaction between age and the neuroanatomical predictors of EF, we hypothesized that:

*Hypothesis 1*. Higher WM integrity (FA) in the forceps minor (FM) and superior longitudinal fasciculus (SLF) would predict better EF performance.
*Hypothesis 2*. Larger GM volumes in a frontal-parietal system would predict better EF performance.
*Hypothesis 3*. The influence of the FM would be more robust as participant age increases, while the predictive value of GM and SLF would be constant across ages.

## Methods

### Participants

More detailed methodology is available in Supplementary Materials. A total of 444 adult participants (Table 1) with complete neurocognitive and scan data were included in these analyses. The data were selected from de-identified phenotypic and neuroimaging data for 645 participants, which was made available via the enhanced Nathan Kline Institute – Rockland Sample (NKI-RS), an open-access, cross sectional, community sample (Nooner, et al., 2012). Rockland County’s economic and ethnic demographics are representative of the United States census (U.S. Census Bureau, 2009), making the NKI-RS generalizable to the U.S. population. Data use agreement was accepted by NKI-RS and data handling procedures were approved by the Institutional Review Board at Suffolk University.

**Table 1.** Demographics for full sample (*n* = 444)

|  |  | *M(SD)* |
| --- | --- | --- |
| Age |  | 48.45(17.33) |
| Education |  | 15.67(2.24) |
|  |  | % |
| Sex |  |  |
|  | Male | 34.9 |
|  | Female | 65.1 |
| Race |  |  |
|  | Native American | 0.7 |
|  | Asian | 4.3 |
|  | Black | 16.4 |
|  | White | 75.7 |
|  | Other | 3 |
| Ethnicity |  |  |
|  | Hispanic | 10.4 |
|  | Non-Hispanic | 89.6 |

Participants were excluded for major psychiatric or neurological conditions, or problems with MRI scans (see Supplementary Material 1. Participant Exclusion). The NKI-RS was selected for this study due to its unique properties. With minimal exclusion criteria, the NKI-RS represented the distribution of psychiatric disorders among the U.S. population. Consistent with epidemiological research showing lifetime prevalence of 46.4% (Kessler, Chiu, Demler, & Walters, 2005), 45.7% of this sample had at least one *relatively less severe* DSM-IV Diagnosis (current or past), as determined by a semi-structured clinical interview. In addition to imaging data and demographic data, the NKI-RS included a battery of neurocognitive tasks that was used to assess brain-behavior relationships.

### Measures

#### EF

In order to examine the unity aspect of EF, it was conceptualized as a latent variable created from five indicators. As an assessment of broad EF, participants completed the Delis-Kaplan Executive Function System Test (D-KEFS; Delis, Kaplan, & Kramer, 2001) and the Penn Computerized Neurocognitive Battery (PennCNB; Gur et al., 2001). Because age was a predictor variable of interest, raw scores were used rather than age-scaled scores. All measures were standardized and timed tasks were reverse coded, so that all positive scores reflected better performance. See Supplementary Table 1 and 2 for descriptive statistics of EF measures, and Supplementary Material for detailed descriptions of neuropsychological assessment.

#### D-KEFS

Measures of EF were extracted from the following subtests: Trails (TMT), Design Fluency (DF), Color Word Interference (CWIT), Tower, and Verbal Fluency (VF). For each measure the raw scores, selected from the array of subtest scores, were selected based on the literature and included in analyses (Gunning-Dixon & Raz, 2000; Kaplan, et al., 2001).

#### PennCNB N-Back Task (N-Back)

This measure assesses working memory (Gur et al., 2001) and has been validated with functional neuroimaging meta-analysis showing activation in front-parietal networks (Owen, McMillan, Laird, & Bullmore, 2005).

#### Imaging

Diffusion weighted images (voxel size = 2×2×2 mm, 137 directions) and structural T1 MPRAGE (voxel size = 1 × 1 × 1 mm) were acquired using a 3.0 T Siemens Trio scanner (see Supplementary Material 3. Imaging). Unless otherwise stated, data were obtained from NKI-RS in their raw form and processing was completed as part of this investigation.

#### GM Image Processing

T1 MPRAGE images were automatically reconstructed in Freesurfer 5.3 (http://surfer.nmr.mgh.harvard.edu; see Supplementary Material 3.1 GM Image Processing). Cortical volumes from the Desikan-Killiany atlas (Desikan et al., 2006) were calculated as a ratio of estimated total intracranial volume (Buckner et al., 2004), and averaged across hemispheres. Volumes from frontal and parietal regions were chosen to reflect a unitary biological construct associated with EF (Collette et al., 2005; Neindam et al., 2012; Spreng, Sepulcre, Turner, Stevens, & Schacter, 2013), and constructed using measurement models for latent variables.

#### WM Image Processing

DTI was processed using TRACULA, included in Freesurfer 5.3 (Yendiki et al., 2011; see Supplementary Material 3.2 WM Image Processing). Fractional anisotropy (FA) from two WM pathways relevant to the EF and the GM latent variable was extracted (Schmahmann, & Pandya, 2009): the FM and parietal branch of the SLF. The FM, which courses through the anterior corpus callosum, is the largest anterior inter-hemispheric tract. The SLF, a major intrahemispheric deep WM tract connects frontal and parietal regions indicated in many EF tasks. The SLF FA values were averaged across hemispheres.

### Statistical Analysis Plan

This study took a structural equation modeling approach (Bentler & Weeks, 1980) to assess both *direct* and age-mediated *indirect* relationships between brain structures (Figure 1, Figure 2, and Figure S1), and EF (the criterion). This approach was chosen to (1) examine the underlying common factor of EF and (2) assess the relative GM and WM contributions. EF and GM were assessed as latent variables given the available array of appropriate variable constituents. FM, SLF, and age were assessed as indicators.

In order to examine the specific effect of compensatory recruitment in older adults, the sample was split at age 55, based on the previously reported age-marker of compensatory dedifferentiation (Rypma & D’Esposito, 2000). We assessed model fit on two nested samples, older adults (age 55-85; *n* = 168) and younger adults (age 20-54; *n* = 276), and compared them to the full sample.

### Structural Equation Modeling

Development of the latent variables and the measurement model occurred in two phases: exploratory and confirmatory (see Supplementary Material 4. Structural Equation Modeling). The EF and GM latent variables were created, and a structural model evaluated the influences of the age, GM, FM, and SLF on EF, as indicated by the standardized regression weights of corresponding pathways at or below statistical threshold (*p* ≤ 0.05). Specifically, the mediating effect of age between different neural substrates and EF was compared. In addition to the model fit indices described for the measurement model, changes in the Akaike information criterion (ΔAIC > 2) was used for model comparison. The use of ΔAIC was appropriate as all variables were included in all models, and could therefore be considered nested.

Bootstrapping was completed to create 95% confidence intervals and significance values for both direct and indirect paths, with 5000 bootstrapped samples extracted (Preacher & Hayes, 2008). The standardized regression weights for total, direct, and indirect effects were used to compare associations at or below statistical threshold (*p* < 0.05)

## Results

### Pre-analysis

Correlations between all indicator variables can be found in Table 2. The final factor fit for the latent constructs of EF (χ^2^ = 22.65, *p* = 0.007) and GM (χ^2^ = 239.53, *p* <0.001) was good (see Supplementary 5. Pre-analysis). The final EF and GM latent variables were derived from the indicators found in Table 3.

**Table 2.** Correlations between all indicator variables.

|  | Age | SLF | FM | POP | PT | POR | MOFC | SF | CMF | RMF | RACC | PCUN | IP | TMT | Tower | CWIT | N-Back |
| --- | --- | --- | --- | --- | --- | --- | --- | --- | --- | --- | --- | --- | --- | --- | --- | --- | --- |
| Age | --- |  |  |  |  |  |  |  |  |  |  |  |  |  |  |  |  |
| SLF | -0.06 | --- |  |  |  |  |  |  |  |  |  |  |  |  |  |  |  |
| FM | -.32** | .40** | --- |  |  |  |  |  |  |  |  |  |  |  |  |  |  |
| POP | -.22** | -0.03 | -0.08 | --- |  |  |  |  |  |  |  |  |  |  |  |  |  |
| PT | -.27** | 0.01 | -0.03 | .67** | --- |  |  |  |  |  |  |  |  |  |  |  |  |
| POR | -.20** | -0.08 | -0.08 | .53** | .59** | --- |  |  |  |  |  |  |  |  |  |  |  |
| MOFC | -.17** | -.10* | -.11* | .59** | .55** | .63** | --- |  |  |  |  |  |  |  |  |  |  |
| SF | -.29** | -0.02 | -0.04 | .69** | .68** | .66** | .70** | --- |  |  |  |  |  |  |  |  |  |
| CMF | -.26** | -0.06 | -0.04 | .57** | .52** | .52** | .55** | .67** | --- |  |  |  |  |  |  |  |  |
| RMF | -.26** | -0.06 | -0.09 | .57** | .62** | .68** | .69** | .79** | .65** | --- |  |  |  |  |  |  |  |
| RACC | -.14** | -0.09 | -0.09 | .48** | .51** | .53** | .56** | .62** | .54** | .63** | --- |  |  |  |  |  |  |
| PCUN | -.23** | -0.01 | -0.07 | .62** | .64** | .65** | .70** | .79** | .63** | .74** | .60** | --- |  |  |  |  |  |
| IP | -.26** | -0.04 | -0.08 | .62** | .62** | .59** | .64** | .73** | .56** | .70** | .59** | .74** | --- |  |  |  |  |
| TMT | -.27** | .15** | .16** | 0.09 | 0.06 | 0.02 | 0.08 | .11* | 0.09 | .10* | 0.08 | 0.06 | 0.08 | --- |  |  |  |
| Tower | -.12* | 0.01 | 0.05 | 0.07 | 0.05 | 0.01 | .10* | 0.08 | 0.05 | .10* | 0.05 | 0.04 | 0.04 | .39** | --- |  |  |
| CWIT | -.22** | .15** | .15** | 0.01 | 0.03 | -0.02 | 0.04 | 0.08 | 0.00 | 0.05 | 0.01 | 0.01 | 0.01 | .53** | .28** | --- |  |
| N-Back | -0.07 | .11* | 0.03 | 0.00 | -0.03 | 0.02 | 0.01 | 0.04 | 0.01 | 0.04 | -0.01 | 0.00 | -0.01 | .40** | .31** | .29** | --- |
| DF | -.17** | .10* | .12* | 0.07 | 0.02 | -0.02 | 0.03 | 0.07 | 0.05 | 0.03 | -0.00 | -0.00 | 0.04 | .40** | .22** | .34** | .19** |

Note. \* $p < .05$ \*\* $p < .01$ \*\*\* $p < .001$ . Pearson's $r$ . Abbreviations are as follows: rostral anterior cingulate (RACC), superior frontal (SF), rostral middle frontal (RMF), caudal middle frontal (CMF), pars opercularis (POP), pars triangularis (PT), pars orbitalis (POR), precuneus (PCUN) and inferior parietal (IP) regional volumes.

### Measurement Model

To understand the association between the constituents, both latent variables and stand-alone indicators, a measurement model was created in which EF and GM were set as related to one another (Supplementary Figure 1). To set the metric of latent variables, the first factor loading of each latent variable was set to 1. Model fit was good (χ^2^ = 353.89, df =228, Cmin/df = 1.55, CFI = 0.98, RMSEA = 0.03). All indicators were significantly associated to their latent variable (*ps* <0.01; factor loadings of above 0.40). There was no significant change in model fit after setting the bidirectional path between the two latent variables. However, the EF and GM latent variables were significantly associated (*r* = 0.11, *p* = 0.016).

**Table 3.** Confirmatory factor loadings of indicators on latent variables.

|  | EF | GM |
| --- | --- | --- |
|  | Factor Loading (95% CI) |  |
| DF | .57(.48-.64) | --- |
| N-Back | .49(.34-.63) | --- |
| CWIT | .65(.54-.74) | --- |
| TMT | .80(.71-.87) | --- |
| Tower | .50(.39-.59) | --- |
| IP | --- | .82(.78-.85) |
| PCUN | --- | .87(.84-.89) |
| RACC | --- | .70(.65-.75) |
| RMF | --- | .86(.83-.89) |
| CMF | --- | .73(.68-.78) |
| SF | --- | .91(.89-.93) |
| POR | --- | .74(.66-.79) |
| PT | --- | .76(.71-.79) |
| POP | --- | .74(.69-.78) |

### Structural Model

Age, FM, and SLF were added into the model as indicators. Model fit remained adequate (χ^2^= 878.95, df =366, Cmin/df = 2.40, CFI = 0.93, RMSEA = 0.04). Directional paths from age, GM, FM, and SLF to EF were specified to assess the influence of each variable on EF, while controlling for the level of the other variables. Additionally, the effect of age as a mediator was considered by specifying directional paths from each structure to age. Variables were entered into the model together, so the effect of each variable was examined while holding the other variables constant. Four different models were compared in the full sample (Figure 1). Provided that all other model fit indices were comparable, the best model was chosen based on ΔAIC. The fit of the best model was then examined in the split age sample to assess the direct and indirect pathways indicated by the dedifferentiation hypothesis.

**Figure 1.**
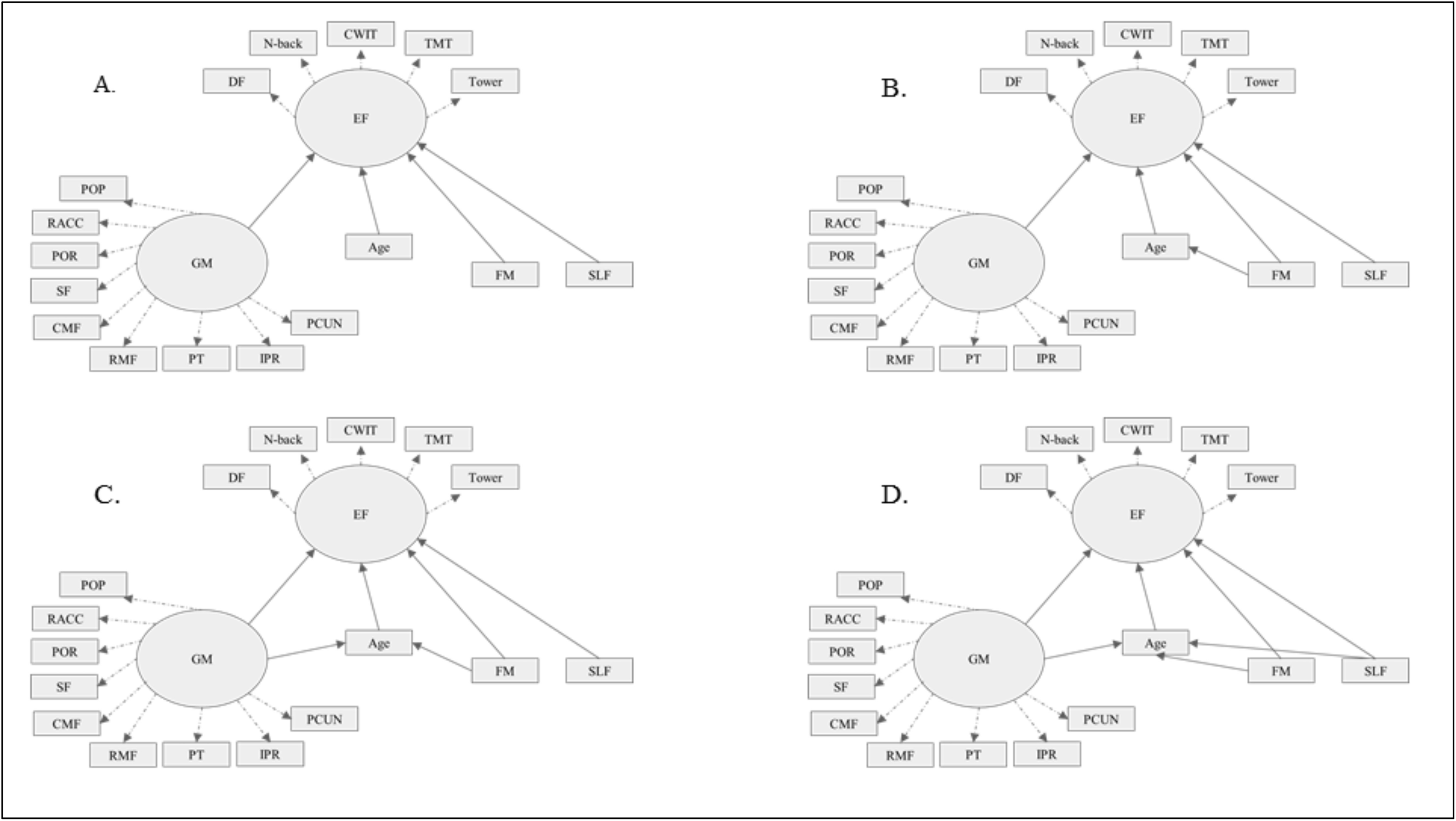
Four structural models that were compared. Model D represents the model with the best fit. Dashed lines indicate factor loadings to latent variables. Abbreviations are as follows: rostral anterior cingulate (RACC), superior frontal (SF), rostral middle frontal (RMF), caudal middle frontal (CMF), pars opercularis (POP), pars triangularis (PT), pars orbitalis (POR), precuneus (PCUN) and inferior parietal (IP) regional volumes. Model fit indices for the four models were as follows: (A) χ^2^ = 797.14, df = 354, *p* < .001, Cmin/df = 2.26, CFI = 0.94, RMSEA = 0.04, AIC = 1109.14; (B) χ^2^ = 788.67, df = 351, *p* < 0.001, Cmin/df = 2.25, CFI = 0.94, RMSEA = 0.04, AIC = 1106.87; (C) χ^2^ = 732.55, df = 348, *p* < .001, Cmin/df = 2.10 CFI = 0.94, RMSEA = 0.04, AIC = 1056.25; (D) χ^2^ = 643.50, df = 345, *p* < 0.001, Cmin/df= 1.87 CFT = 0.96 RMSEA = 0.03 ATC = 973.50

### Full Sample

For the full sample (age 20-85, *n*= 444), the best model by ΔAIC had good fit (χ^2^ = 643.50, df = 345, *p* < 0.001, Cmin/df = 1.87, CFI = 0.96, RMSEA = 0.03). The effect of GM (*β* = 0.13, *p* = 0.003) and FM (*β* = 0.17, *p*= 0.008) on EF was mediated by age (Figure 2). The slopes from the model indicated a 690 mm^3^ increase in GM volume was associated with a 0.09 *SD* increase in EF performance, and a 0.06 increase in FM FA, was associated with a 0.08 *SD* increase in EF performance. For FM and GM, there was full mediation (Baron & Kenny, 1986), with the indirect, age-mediated pathway accounting for 67.3% and 74.8% respectively of the brain structure-EF relationship. CIs and p-values generated from bootstrapped samples (Table 4) for the standardized regression weights of the indirect effect supported this interpretation, for both GM (*β* = 0.10, *p* < 0.001) and FM (*β* = 0.11, *p* < 0.001).

The direct path from SLF (*β*= 0.14, *p =* 0.023) to EF was significant, while the age-mediated indirect path accounted for only 21.2% of the brain structure-EF relationship (Figure 2) and was not significant (*β*= −.02, *p* > 0.05). The model slope indicated a 0.05 increase in SLF FA was associated with a 0.07 *SD* increase in EF performance. This suggests that age played an insubstantial role in mediating the relationship between EF and SLF.

**Figure 2.**
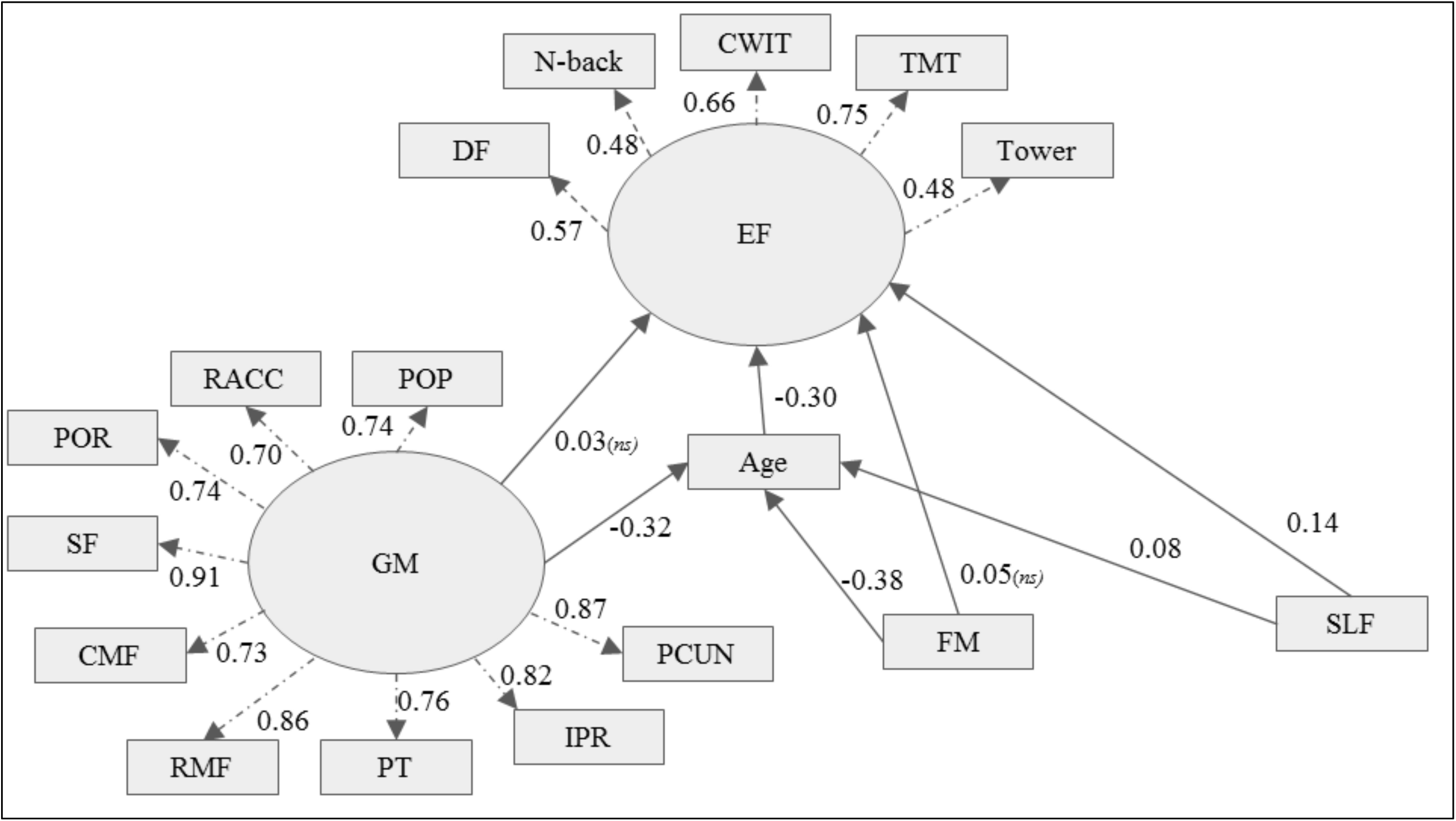
Final model for the full sample (*n* = 444). Dashed lines indicate factor loadings to latent variables. Abbreviations are as follows: rostral anterior cingulate (RACC), superior frontal (SF), rostral middle frontal (RMF), caudal middle frontal (CMF), pars opercularis (POP), pars triangularis (PT), pars orbitalis (POR), precuneus (PCUN) and inferior parietal (IP) regional volumes. Unless otherwise indicated as not significant (ns), all pathways are significant.

### Older Subsample

Group statistics for indicator variables can be found in Supplementary Table 3. Model fit in the older adults (*n* = 168) was comparable to that of the full sample (χ^2^ = 159.66, df = 115, *p* = 0.004, Cmin/df = 1.39, CFI = 0.96, RMSEA = 0.05). However, the RMSEA CI was wider in the older group (range = 0.04) compared to the full sample (range = 0.01), which suggested lower precision and less certainty of model fit (Kenny, Kaniskan, & McCoach, 2015). However, the CI was still sufficiently narrow to infer accurate estimation of relationships (Byrne, 2013). This was also reflected in the CIs and p-values for the standardized total, direct, and indirect relationships between EF GM and SLF (Table 4).

The age-mediated pathway from FM to age to EF was significant for older adults only (*β* = 0.06, *p* = 0.016) with the indirect path accounting for 47.4% of the relationship (Table 4). Comparatively, the age-mediated pathway for GM accounted for only 6.3% of the relationship. Although the direct pathway from SLF to EF appeared strongest (*β* = 0.19, *p* = 0.055), the corresponding CI included zero (−.01. 0.36) and it was not significant. This could be an artifact of the lowered precision of the model (and thus wider confidence intervals). The direct and indirect paths from GM and SLF to EF were not significant in the older adults.

Younger Subsample. For younger adults, the model fit indices ranged from good to acceptable (χ^2^ = 216.77, df = 115, *p* < 0.001, Cmin/df = 1.89, CFI = 0.95, RMSEA = 0.06). The relationships between EF and neuroanatomical structures were not significant (Table 4). Possibly due to restricted variance in EF among younger adults (0.57) compared to older adults (1.01), age was no longer associated with EF (Beta = −.09, p = 0.253) and all age-mediated pathways became non-significant.

**Table 4.** Standardized total, direct, and indirect effects (*β*) of brain structures on executive functioning.

|  | Full (Age 20-85) |  |  | Younger (Age 20-54) |  |  | Older (Age 55-85) |  |  |
| --- | --- | --- | --- | --- | --- | --- | --- | --- | --- |
|  | GM | FM | SLF | GM | FM | SLF | GM | FM | SLF |
| Total | .13** | .17** | .12 | .05 | .09 | .14 | .11 | .14 | .15 |
| Direct | .03 | .05 | .14* | .02 | .06 | .14 | .10 | .07 | .19 |
| Indirect | .10*** | .11*** | -.02 | .02 | .03 | .00 | .01 | .06* | -.04 |

*Note.* \* $p < .05$ \*\* $p < .01$ \*\*\* $p < .001$

## Discussion

The principal findings of our study were that: (1) while higher FA in both the FM and the SLF was associated with better EF performance, only FM was fully mediated through age (2) the effect of GM on EF was similarly mediated through age, and (3) the influence of FM is more salient for older adults compared to SLF and frontal-parietal GM as evidenced by significance of that pathway only among older adults.

Compared to frontal GM and the intrahemispheric WM pathway (SLF), the interhemispheric WM pathway (FM) explained the most variance in EF for older adults, consistent with the dedifferentiation hypothesis of compensatory homologous recruitment (Cabeza et al, 2002). The consistency of this relationship after the age-split suggests that the FM is a particularly relevant substrate for high performing, community dwelling older adults.

The relationship between intrahemispheric WM and EF was not mediated by age, but instead had a direct effect on EF in the full age range. This suggests that intrahemispheric structures necessary for recruitment in the parietal region are relevant across the lifespan, not as an age-related compensatory mechanism. Although many neuropsychological tests are designed as “frontal batteries”, tasks of EF often require recruitment in multiple regions across a frontal-parietal network (Collette et al., 2005). The direct effect of intrahemispheric WM on EF suggests that the recruitment of intrahemispheric regions may stay relatively constant across the lifespan, reflecting the centrality of frontal-parietal recruitment is central to these tasks. The increased variability between intrahemispheric WM and EF in the older adults also supports the conclusion that intrahemispheric recruitment is not the primary mechanism of age-based compensation in community dwelling older adults.

The influence of GM on EF was mediated by age, which is not consistent with literature showing the static association between single EF measures and brain regions (Yuan & Raz, 2014). Our findings may be a consequence of reduced variance in younger adults (age 20-54) compared to older adults (age 55-85). The reduced variance seems to be the result of ceiling effects in the timed tasks used in the latent variable. Notably though, the relationship between GM and EF was also significantly reduced in older adults, as compared to the full sample. This may be because the network is of paramount importance, not its constituent parts. It is also possible that influence of decreasing volume in GM regions may be more salient in middle aged adults, which were not captured as an age group due to limits in sample size.

Future research should investigate specific thresholds for age-mediated pathways between brain structures and EF, specifically the changing effects of GM in the transition period of middle age. We would also expect that these relationships would change in the presence of neuropathology. While the NKI-RS offers unique advantages as a highly representative sample, these results may not generalize to certain clinical populations characterized by pathological and age-related structural changes. In clinical samples there may be additional recruitment needs both inter- and intrahemispherically, for which identification is critical in dementia progression prevention.

These results suggest that the neural substrates of EF are not static across the lifespan, but change in later life, as paralleled in early lifespan work (Horowitz-Kraus, Holland, & Freund, 2016). The impact of these findings, in terms of variance accounted for, was moderately small, but does have theoretical and clinical ramifications. As detection and prevention programs for cognitive decline identify neural targets of interest, WM substrates may be more relevant for geriatric populations. Consistent with previous work in dedifferentiation, the efficiency of activation mediated through WM may be more important than any one GM region (i.e. salience of network over node).

## Acknowledgements

The authors would like to acknowledge the following people and organizations for their contributions:

The NKI-Rockland Sample Initiative for providing the data used in these analyses (data collection funded through NIMH BRAINS R01MH094639-01).

The Suffolk University Psychology Department for their support of doctoral students and David Gansler’s Lab, and the contributions of graduate student Mrs. Sarah Levy.

